# Chemotherapy-induced transfer of apoptotic machinery in extracellular vesicles between somatic and germ cells of the testis: mechanistic insights into onco-fertility preservation in pre-pubertal boys

**DOI:** 10.64898/2026.01.17.699063

**Authors:** Michael P Rimmer, Pamela Holland, Daniel C Rabe, Shannon L Stott, Christopher D Gregory, Rod T Mitchell

## Abstract

Extracellular vesicles (EVs) are increasingly recognized as critical mediators of intercellular communication, not least during cellular stress or therapy. While EV signalling is well-studied in various tissues, its role in the prepubertal testicular environment is not well understood. Chemotherapy, commonly used in paediatric oncology, poses a significant risk to spermatogonial stem cells (SSCs) and may affect long-term fertility in cancer survivors. The role of EVs in chemotherapy-induced testicular damage in these patients is unknown and may be important for developing new fertility preservation methods.

Immortalised murine Sertoli (TM4) and spermatogonial (GC1-spg) cell lines were used to investigate cisplatin-induced changes in EV biogenesis, release, and function in an in vitro model of the prepubertal testicular microenvironment. Our findings indicate that cisplatin significantly increases EV secretion and internalisation by recipient cells. Notably, EVs from cisplatin-exposed Sertoli cells exhibit a novel pro-apoptotic phenotype when co-cultured with chemotherapy-naïve Sertoli cells. Proteomic profiling of these EVs shows enrichment of apoptosis-regulatory proteins including caspases, activating Caspase-3/7 in recipient Sertoli cells. Conversely, germ cells exposed to Sertoli cell-derived EVs displayed reduced levels of apoptosis as well as a chemoprotective role to germ cells undergoing treatment with cisplatin.

These findings indicate a dual role for Sertoli cell-derived EVs in mediating (1) apoptosis in Sertoli cells and (2) protection of germ cells following cisplatin exposure. The presence of pro-apoptotic molecules, especially caspases, in chemotherapy-induced Sertoli cell EVs provides mechanism for the induction of somatic cell apoptosis. Furthermore, their protective effects on germ cells demonstrate the complexity of EV-mediated signalling between testicular cell types. Manipulating EV biogenesis and cargo loading could be a promising approach to reduce chemotherapy-related gonadotoxicity and preserve fertility in childhood cancer patients.

## 1 Introduction

Extracellular vesicles (EVs) are small lipid-delimited structures released from cells into the extracellular milieu that mediate cell-to-cell communication, influencing recipient cell activity both locally and at distant sites. This occurs through mechanisms including cell surface receptor signalling, cargo release at the cell surface, or internalization and transport to specific subcellular loci (Joshi et al., 2020, O’Brien et al., 2022). EVs have been well-characterized in numerous biological and clinical contexts (Thery et al., 2002, Zhang et al., 2021, Yates et al., 2022).

EV cargoes, including RNAs, DNA, lipids, proteins, and metabolites, are incorporated into the recipient cell based on the parent cell state (Tucher et al., 2018, Maas et al., 2017). Differential cargo expression has led to their increasing use as biomarkers for diagnosis and prognosis in various pathologies (Rimmer et al., 2021a, Gregory and Rimmer, 2023, He et al., 2021, Leggio et al., 2021). While much research focuses on EV roles in homeostasis (Yates et al., 2022), there is increasing focus on EVs released by cells following disease transformation or in response to treatment (e.g., chemotherapy).

Fertility in males is dependent on differentiation of spermatogonial stem cells (SSC) into sperm, which requires cell-cell communication between somatic cells and their neighbouring germ cells. Sertoli cells, located in close contact with the germ cells in the seminiferous tubules, represent the key somatic cell population supporting SSCs in the prepubertal testis and spermatogenesis in adulthood. Testicular SSCs are sensitive to gonadotoxic insult, notably chemotherapeutics, which can cause infertility (Chow et al., 2016). Germ cell damage occurs through direct effects on SSCs or indirectly via the supporting somatic cells that contribute to the SSC niche. While sperm banking offers adult men the opportunity to preserve their fertility prior to undergoing gonadotoxic therapy, this is not an option for prepubertal boys who lack sperm production. Protecting the SSCs from gonadotoxic compounds in this population is a significant unmet clinical need. Understanding germ-soma interactions mediating chemotherapy-induced testicular damage is crucial for identifying fertility preservation strategies.

EVs play numerous key roles in the adult male reproductive tract, including modulating soma-to-germ line communication, with the potential to alter offspring phenotype in mice (Chan et al., 2020) or promoting sperm motility in humans (Frenette et al., 2005). EVs can also facilitate zona pellucida binding and the acrosome reaction between sperm and the oocyte (Frenette and Sullivan, 2001), and enhance fertilisation and capacitation across species (Harayama et al., 1999, Nixon et al., 2019, Sharma et al., 2016, Belleannee et al., 2013, D’Amours et al., 2012, Griffiths et al., 2008, Caballero et al., 2012, Chabory et al., 2009, Tan et al., 2022, Rimmer et al., 2021b). However, the role of EVs in maintaining the SSC niche, supporting germ cell development, and their response to chemotherapy exposure remains unknown.

Cisplatin is an alkylating-like chemotherapeutic used in the treatment of several types of childhood cancer (Tian En et al., 2020). Cisplatin exposure is known to result in gonadotoxicity, loss of SSCs and risk of future infertility (Chow et al., 2016, Matilionyte et al., 2022, Tharmalingam et al., 2020). We investigated how cisplatin may alter EV release from prepubertal Sertoli cells and the impact of those EVs on other key cell populations. Intriguingly, chemotherapy treatment led to the production of EVs with a pro-apoptotic phenotype, enriched with caspases, which induce apoptosis in treatment-naive Sertoli cells, following incubation. By contrast, Sertoli cell-derived EVs, including those secreted under control conditions, were found to exert a chemoprotective effect against cisplatin-induced toxicity.

## 2 Materials and Methods

### 2.1 Cell culture

The immortalised mouse cell lines GC1-spg, representing Type-B spermatogonia, and TM4, representing Sertoli cells, were used in this work. GC1-spg cells were created using isolated germ cells at post-natal day 10 (Hofmann et al., 1992) and cultured in DMEM with 10% fetal bovine serum (FBS), while TM4 cells were derived at post-natal day 11-13 (Mather, 1980) and cultured in DMEM:F12 with 5% horse serum (HS) and 2.5% FBS. All cells were cultured at 37 °C and 5% CO_2_.

### 2.2 Production of EV depleted media

EV depletion from FBS and HS was performed by centrifuging whole serum for one hour in Amicon ultra-15 centrifugal filter units (Merck, Germany) with a 3 kDa cut-off for FBS, or a 10 kDa cut-off followed by a 3 kDa cut-off for horse serum, similar to the protocol described here by (Kornilov et al., 2018). EV depletion was confirmed using Nanoparticle Tracking Analysis (NTA) prior to use in cell culture, as described below, to confirm depletion of EVs from serum.

### 2.3 Cisplatin treatment to cells

GC1-spg and TM4 cells were treated with cis-Diamminedichloroplatinum(II) (cisplatin; Sigma-Aldrich, Germany) or water (vehicle control) to produce cisplatin-induced or control EVs. The cisplatin concentrations used ranged from 0.0 μM – 200 μM, reflecting serum levels observed in children undergoing platinum-based chemotherapy (Rajkumar et al., 2016, Veal et al., 2007, Peng et al., 1997). These doses encompass the full dose-response curve for GC1-spg and TM4 cell survival/death.

### 2.4 Cell death quantification using Trypan Blue

GC1-spg and TM4 cells were cultured in EV-depleted media for 24 hours and subsequently exposed to cisplatin at concentrations ranging from 0 to 200 μM for another 24 hours. Cell viability was then assessed using Trypan Blue staining (Gibco, USA), with non-viable cells identified by blue staining and counted accordingly using a haemocytometer with a minimum of four squares counted. A dose-response curve was generated to determine the proportion of non-viable cells at each cisplatin concentration after 24 hours of exposure.

### 2.5 Isolation of EVs from cell culture media

EVs were isolated from cell culture media by first concentrating the media to a final volume of 1 mL using Amicon Ultra-15 centrifugal filter units with a 3 kDa cutoff. The concentrated media containing EVs was then subjected to size-exclusion chromatography using Sepharose CL-6B cross-linked beads (Merk, Germany), with elution performed using Hanks’ Balanced Salt Solution (Corning, USA). To concentrate the EVs further, the EV-containing fractions were further concentrated using the same Amicon Ultra-15 filters.

### 2.6 Nanoparticle tracking analysis

Quantification of EV number and size was undertaken using a Nanosight LM10 instrument (Malvern, UK), capable of reliably detecting EVs up to 1000 μM in size. A minimum volume of 1 mL was analysed by manually introducing the sample into the sample chamber and a 30 second video recorded using cMOS Hamamatsu-60 Orca Flash 2.8 camera and then processed using Nanosight software 2.3. Videos were recorded using a camera level of 15 and analysed using an upper imaging threshold of 15,080 and a detection threshold of 15 with a minimum expected size of 30 nm. Videos were recorded in triplicate and the concentration of EVs present was calculated by taking the mean concentration of all recorded videos. Prior to use, the Nanosight was calibrated with 100 nm latex beads (Nanosphere ^TM^, Thermo Fisher Scientific, USA).

### 2.7 Transmission electron microscopy

EV preparations were diluted to concentrations of 1:10, 1:25, and 1:50 using 1× phosphate-buffered saline (PBS). Subsequently, 10 μL droplets of each dilution were placed onto 200-mesh Formvar/carbon-coated nickel grids (Sigma-Aldrich, Germany). The droplets were blotted to remove excess liquid, then contrast-stained with a 2% methyl cellulose (tylose) and uranyl acetate solution before air drying. The samples were examined using a JEOL transmission electron microscope operated at 80 kV. Images were captured with an AMT digital imaging system equipped with proprietary image capture software (Advanced Microscopy Techniques, Danvers, USA).

### 2.8 Western Blotting

EVs from GC1-spg and TM4 cells were isolated by ultracentrifugation at 100,000 G for 2 hours using a TH-614 swing-out rotor (Sorvall, USA). Following centrifugation, the media was carefully aspirated and discarded without disturbing the EV pellet. The pellet was then lysed on ice for 30 minutes in 100 μL of lysis buffer, consisting of 1X RIPA buffer, 5 μL of Roche cOmplete protease inhibitor (Sigma-Aldrich, Germany), and 1 μL of each phosphatase inhibitor cocktail (Sigma; Cat No: P5726 and P0044). After lysis, the samples were centrifuged at 5,000 rpm for 5 minutes at 4°C, and the clear supernatant was transferred to fresh tubes on ice.

EV lysate (20 μg) was prepared by resuspending in 50 μL of Laemmli sample buffer (Bio-Rad, USA) containing 5% β-mercaptoethanol (Pan Reac, USA). The samples were heated at 70°C for 5 minutes and then loaded onto pre-cast 4-12% Bis-Tris Plus gels (Thermo Fisher Scientific, USA), alongside a pre-stained protein ladder (Thermo Fisher Scientific, USA). Electrophoresis was conducted in NuPAGE MOPS SDS (20x) running buffer (Novex, USA) by running the samples at 200 V for 50 minutes on ice.

Proteins were transferred onto a nitrocellulose membrane (Immobilon Membrane, Thermo Fisher Scientific, USA) by first immersing the membrane in 100% methanol for 10 minutes. The membrane was then incubated in 1x transfer buffer (Thermo Fisher Scientific, USA) for 30 minutes. A semi-dry transfer was performed using a Pierce G2 fast blotter (Thermo Fisher Scientific, USA) set to 25 V and 1.3 A for 7 minutes. After the transfer, the membrane was washed three times in PBS containing 0.1% Tween 20, each wash lasting 5 minutes. Finally, the membrane was blocked in a 1:1 mixture of intercept blocking buffer (Li-Cor, USA) and PBS for 1 hour.

Membranes were probed overnight at 4°C with TSG-101 and alpha-tubulin antibodies (Supplementary Table 1) in a 1:1 mixture of blocking buffer and PBS. Following incubation, the membranes were washed three times for 5 minutes each with PBS containing 0.1% Tween 20 to remove unbound antibodies. Subsequently, the membranes were incubated with secondary antibodies (Supplementary Table 1) for 1 hour at room temperature. After another three washes of 5 minutes each in PBS with 0.1% Tween 20, the membranes were imaged using an Odyssey® FX Imager (LiCor, USA), capturing signals in the 800 nm and 680 nm channels.

### 2.9 dSTORM single EV imaging

EV preparations were stained using an Oxford NanoImager (ONI) EV profile Kit (ONI, UK). EVs (9 μL) were incubated overnight at 4°C with primary fluorescent antibodies (anti-CD9-ATTO-488, anti-CD63-Cy3-568, or anti-CD8-AlexaFluor-647; ONI, UK). Following incubation, each lane of the ONI capture chip was sequentially incubated with 20 μL of ONI surface solution-3 (S3), washed with 100 μL of ONI wash buffer-1 (W1), incubated with 10 μL of ONI surface solution-4 (S4), and finally flushed with 50 μL of W1. All reagents were supplied by ONI, UK.

EVs were resuspended to a final volume of 10 µL using ONI Capture Buffer-1 (C1). The suspension was then loaded into the lanes of the ONI capture chip, washed with 100 µL of ONI Wash Buffer-1 (W1), and fixed with 50 µL of ONI Fixation Solution-1 (F1).

ONI EV capture chips were imaged using an Oxford NanoImager equipped with NimOS software, with image analysis performed on the CODI platform (available at: https://alto.codi.bio/). Prior to imaging, the ONI imaging chamber was maintained at 37°C. Channel alignment across the 488, 660, and 640 channels was verified using TetraSpec microspheres 0.1 μm (Thermo Fisher Scientific, USA), placed in an Ibidi 8-well glass-bottom chamber slide (Ibidi, Germany).

Image acquisition was undertaken in two sequential steps as the ONI utilizes a 640 dichroic split, allowing images to be acquired simultaneously in the 647nm channel (red channel) and either the 488nm or 560nm channel (green channel) and images exported into ONI’s digital platform analysis software CODI and corrected for drift using the platform software.

### 2.10 EV tracking within cells

EVs released by GC1-spg cells were fluorescently labelled with Alexa Fluor 647-NHS Ester by incubating at a 1:1000 dilution at room temperature for one hour (Thermo Fisher Scientific, USA). Excess dye was removed by size exclusion chromatography, and the number of eluted EVs was quantified using NTA. The fluorescently labelled EVs were then incubated with treatment-naïve GC1-spg cells cultured on an Ibidi 8-well glass bottom chamber slide (Cat No: 80807) for an additional 24 hours. EV movement within the cells was monitored at 24 hours, with imaging performed using the 647 nm channel on the ONI system.

Single cells were imaged using the multiple acquisition software of the ONI NimOS, 2000 frames per cell were captured, each frame for 100 ms. NimOS EV tracking analysis software was used to determine the number of individual EVs within each cell that were motile and the speed of each EV.

### 2.11 Quantification of EV uptake by recipient cells

EV preparations were fluorescently labelled with Alexa Fluor 488-NHS Ester at a dilution of 1:1000 (Thermo Fisher Scientific, USA). Excess dye was removed through size exclusion chromatography, and the final EV concentration was determined using NTA. The labelled EVs were incubated with treatment-naïve cells, seeded at 2,500 cells / well in 96-well plates and cultured for 24 hours, and then imaged using the IncuCyte Live Cell Analysis System (Sartorius, Germany). Images were captured every 3 hours over a 24 hour period to monitor the uptake of EVs by cells, identified by their fluorescence at 488 nm.

Using the Incucyte ZOOM software (Sartorius, Germany), images from different time points over the 24 hours period were utilised to create a training library of images. Applying machine learning, the Incucyte ZOOM software was trained to identify a cell using parameters such as minimum and maximum area, object eccentricity (Wurster et al., 2019), and green object count per mm² with the 488 nm channel. Once the reliable identification of 488^+ve^ (EV-associated) cells was confirmed through the training library images and manual inspection, this was used to quantify the amount of fluorescent EV (control or cisplatin-induced) uptake by treatment-naïve cells.

### 2.12 Fluorescent quantification of apoptosis in treatment-naïve cells incubated with cisplatin-induced EVs

EVs (1×10^8^) were generated from TM4 cells treated with varying concentrations of cisplatin (0-200 μM). These cisplatin-induced EVs were purified using size exclusion chromatography as described previously. The EVs were then isolated and quantified by NTA. Treatment-naïve GC1-spg and TM4 cells were seeded at 2,500 cells per well in 96-well plates and cultured for 24 hours. Subsequently, these cells were exposed to the cisplatin-induced EVs for an additional 24 hours. To assess apoptosis, the NucView 488 caspase-3 substrate (diluted 1:1000; Biotium, USA) was added to each well. Staurosporine (20 μM) was used as a positive control for apoptosis.

Wells were imaged using the Incucyte Live Cell Analysis System (Sartorius, Germany). Images were captured every 3 hours over a total period of 24 hours to identify cells, including 488^+ve^ cells that had undergone apoptosis. Similar to the detection of fluorescent EV uptake, a library of training images was developed, and a machine learning algorithm was used to ensure accurate identification of apoptotic cells. The number of green objects per mm^²^ was quantified and exported for further analysis. Apoptosis levels in treatment-naïve cells were compared between those incubated with control EVs and those exposed to EVs released by cisplatin-treated cells.

### 2.13 Fluorescent quantification of apoptosis in cells incubated with cisplatin-induced EVs followed by cisplatin exposure

GC1-spg and TM4 cells were seeded at 2,500 cells / well in 96 well plates and cultured for 24 hours. Cells were then incubated with TM4 EVs, isolated as described above for 24 hours. Subsequently, media and EVs were removed and cells washed and then treated with cisplatin for 24 hours. The impact of cisplatin on GC1-spg cells incubated with EVs was quantified using NucView 488 caspase-3 substrate (1:1000; Biotium, USA) to identify apoptotic cells. A positive control for apoptosis of Staurosporine (5 μM) was also used.

### 2.14 Reversal of apoptosis

To inhibit caspase machinery within EVs, TM4 cells were cultured and EVs isolated and concentrated as outlined above. Prior to undergoing size exclusion chromatography, an additional step was included in which EVs were incubated with 50 μM of the pan-caspase inhibitor Z-VAD(OH)-FMK (zVAD) for 60 minutes at room temperature. To remove cisplatin, cell debris and excess zVAD, EVs were then passed through SEC columns to elute a homogeneous population of EVs.

### 2.15 Proteomics

Five paired samples of EVs released by TM4 cells treated with either 10 μM cisplatin or control media were assessed for their protein cargoes. EV preparations were concentrated by ultracentrifugation and pelleted EVs were then lysed in 25 μL of lysis buffer.

5μg of protein was digested for mass spectrometry (MS) using the Filter Aided Sample Preparation (FASP) protocol, as previously described (Wiśniewski et al., 2009). Proteins were diluted in 150 μL of denaturation buffer containing 8M Urea in 50 mM ammonium bicarbonate (ABC) and placed on top of an Amicon ultra-30 centrifugal filter unit with cut-off of 30 kDa (Sigma Aldrich, Germany). Samples were spun at 14,000 x g for 20 minutes, washed with 200 μL of denaturation buffer followed by further centrifugation under the same conditions. The protein samples were then reduced by the addition of 100 μL of 10 mM dithiothreitol (Sigma Aldrich, Germany) in denaturation buffer for 30 minutes at ambient temperature and alkylated by adding 100 μL of 55 mM iodoacetamide (Sigma Aldrich, Germany) in denaturation buffer for 20 minutes at ambient temperature in the dark. Two washes with 100μl of denaturation buffer and two with digestion buffer (50 mM ABC) were performed before the addition of trypsin (Pierce, USA). The ratio of protease to protein was 1:50 and proteins were digested overnight at 37°C. Following digestion, samples were spun at 14,000 x g for 20 minutes and the flow-through containing digested peptides was collected. Filters were then washed one more time with 100μl of digestion buffer and the flow-through was collected again. The eluates from the filter units were acidified using 20 μl of 10% Trifluoroacetic Acid (TFA) (Sigma Aldrich, Germany) and spun onto Stage Tips (Thermo Fisher Scientific, USA) prior to assessment using MS (Rappsilber et al., 2007). Peptides were eluted in 40 μL of 80% acetonitrile in 0.1% TFA and concentrated down using vacuum centrifugation (Concentrator 5301, Eppendorf, UK). The peptide sample was then prepared for LC-MS/MS (liquid chromatography with tandem mass spectrometry) analysis by diluting it to 5 μL in 0.1% TFA.

LC-MS analyses were undertaken using an Orbitrap Exploris™ 480 Mass Spectrometer (Thermo Fisher Scientific, USA), coupled on-line to an Ultimate 3000 HPLC (high performance liquid chromatography; Dionex, Thermo Fisher Scientific, USA). Survey scans were recorded at 120,000 resolution (scan range 350-1650 m/z) with an ion target of 5.0e6, and injection time of 20 ms. MS2 Data Independent Acquisition was performed in the Orbitrap Mass Spectrometer (Thermo Fisher Scientific, USA) at 30,000 resolution with a scan range of 200-2000 m/z, maximum injection time of 55 ms and AGC target of 3.0E6 ions. The inclusion mass list with the corresponding isolation windows containing empirically labelled peptides and their charge enabling peptide identification are shown in Supplementary Table 2.

The DIA-NN software platform version 1.8.1 (Demichev, 2020) was used to process the raw files and a protein search was conducted against the Mus musculus Uniprot database, a repository of data combining UniProt Knowledgebase, UniProt Reference Clusters and UniProt Archive (released in July, 2017 and available at: https://www.uniprot.org/). The parameters for peptide length range, precursor charge range, precursor m/z range and fragment ion m/z range as well as other software parameters were used with their default values.

### 2.16 Gene ontology and search tool for the retrieval of interacting genes and proteins

Ontology of molecular functions and biological functions was undertaken. Gene ontology of identified EV proteins was analysed using the open-source software PANTHER (available at: http://pantherdb.org/). Search tool for the retrieval of interacting genes and proteins (STRING) (String Consortium 2022, available at: https://string-db.org/) analysis was undertaken of proteins identified with functions in apoptosis and reproduction. Data were represented visually with known interactions between proteins and pathway function denoted by a line between proteins on a Kyoto Encyclopaedia of Genes and Genomes pathways map.

### 2.17 Statistical analysis

All statistical analysis was undertaken using GraphPad PRISM, version 8 (GraphPad Software Inc., USA). When assessing the differences in the proportion of fluorescent EV uptake in treatment-naïve GC1-spg or TM4 cells, a 1-way ANOVA was conducted. The fold change in EV uptake was determined by comparing the amount of EVs taken up by cells exposed to EVs from cisplatin-treated cells versus those from control-treated cells. Data are represented as mean ± standard deviation with statistical significance set at p<0.05.

Apoptosis was investigated using two platforms, the Incucyte Zoom and CellCyteX. To assess differences in levels of apoptosis in GC1-spg or TM4 cells incubated with EVs, a 1-way ANOVA was conducted. The fold change in apoptosis was determined by comparing the number of apoptotic cells incubated with EVs released by cisplatin-treated cells, to the number of apoptotic cells incubated with control EVs (those released by cells treated with control media). Data are represented as mean ± standard deviation with statistical significance set at p<0.05.

To gain more detailed insights into our apoptosis assays, we employed a second imaging platform, the CellCyteX, which provides higher resolution than the Incucyte Zoom platform. We assessed differences in apoptosis levels in TM4 cells incubated with EVs derived from cells treated with cisplatin or control media and then incubated with zVAD or PBS. A one-way ANOVA was used to analyse these data. We compared the number of apoptotic cells per mm², incubated with EVs from different sources, including EVs from cisplatin-treated cells and control EVs, with or without zVAD incubation, to controls. Data are represented as mean ± standard deviation, with statistical significance set at p < 0.05.

For proteomics analysis, data were exported to Microsoft Excel and analysed in Perseus software, version 1.6.2.1 (MaxQuant, available at: https://maxquant.net/perseus/; Tyanova et al., 2016) and GraphPad PRISM, version 8 (GraphPad Software Inc. USA). Raw values were log transformed for data analysis. Missing value imputation was undertaken using Perseus software based on the normal distribution of the existing values. A p value of Log < 0.05 was used when undertaking multiple paired t tests in GraphPad PRISM. Data were represented in a volcano plot as protein value in EVs released by cells treated with control media, minus, protein values in EVs released by cells treated with 10μM cisplatin. Differential expression of proteins was considered significant if the Log p value was > 1.3 (p=0.05).

## 3 Results

### 3.1 GC1-spg germ cells and TM4 Sertoli cells produce EVs

We began by characterising the EVs released by GC1-spg and TM4 cells. For the conditions investigated, GC1-spg and TM4 EV size assessment using NTA demonstrated that the mean size of GC1-spg EVs was 200 nm and that of TM4 EVs was 250 nm (Fig. 1A, B), within the detection limits of the Nanosight. Western Blotting for the EV marker TSG-101 revealed that the bulk preparations of GC1-spg and TM4 EVs both contained TSG-101 (Fig. 1C, D), while individual EV dSTORM imaging for CD9, CD63, and CD81 showed numerous EVs to be positive for at least one of these markers (Fig. 1E-J). TEM analysis of GC1-spg and TM4 EVs displayed the classical rounded appearance with an average size of < 200 nm (Fig. 1K-V).

**Figure 1:**
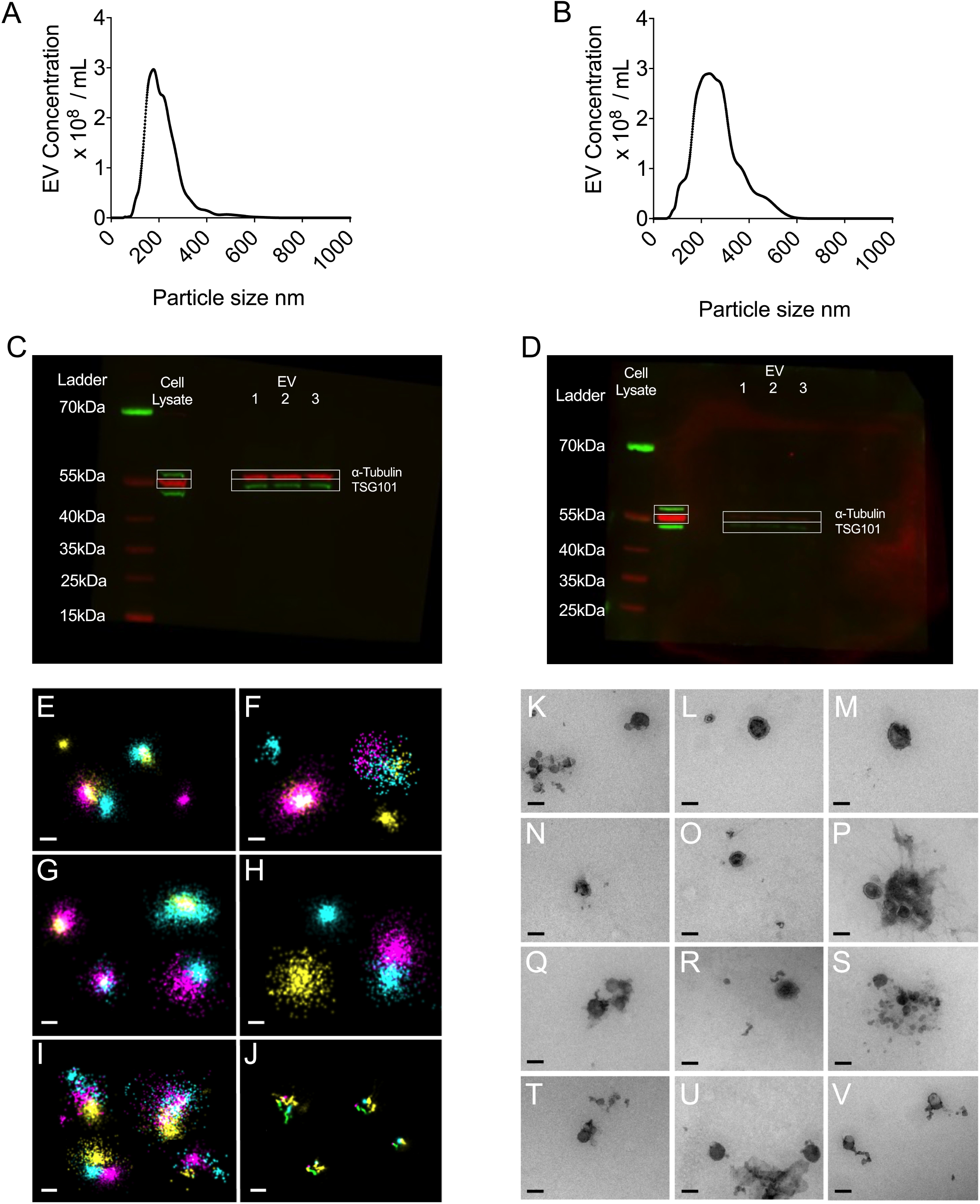
Characterisation of EVs: Nanoparticle tracking analysis (NTA) showing the size distribution of EVs isolated from A) GC1-spg cells and B) TM4 cells cultured in control media. Western blot of EV lysate released by C) GC1-spg and D) TM4 cells cultured in control media forTSG-101 (45kDa, green) and alpha-tubulin control (55kDa, red), with n=3. Representative dSTORM images of EVs released by GC1-spg cells treated with E) control media or F) 10 µm cisplatin, TM4 cells treated with G) control media or H) 10 µm cisplatin, and I) Oxford NanoImager control EVs and J) TetraSpecTM Microspheres stained for CD9-cyan, CD63-yellow, and CD81-magenta. Representative transmission electron microscopy images of EVs released by GC1-spg cells cultured in control media (K,L,M) and 10 µm cisplatin (N,O,P), as well as EVs from TM4 cells cultured in control media (Q,R,S) and 10 µm cisplatin (T,U,V). All scale bars -100 nm.

### 3.2 Cisplatin exposure increases EV release from TM4 and GC1-spg cells and increases EV uptake in chemotherapy-naïve cells

Exposure of GC1-spg and TM4 cells to a range of cisplatin doses (0-200 μM) resulted in cell death varying from 0% to 90% (Supplementary Fig. 1A, 1B). EV release from cisplatin-treated GC1 cells was significantly increased at 10 µM (7.2 x 10^8^ / mL vs 13.6 x 10^8^ / mL, p=0.009) and 100 µM (7.2 x 10^8^ / mL vs 16.3 x 10^8^ / mL, p=0.0002) concentrations, compared to vehicle-exposed controls (Fig. 2A), as well as TM4 cisplatin-treated cells at 10 µM (2.8 x 10^8^ / mL vs 6.4 x 10^8^ / mL, p<0.0001) and 100 µM (2.8 x 10^8^ / mL vs 5.0 x 10^8^ / mL, p=0.0015) concentrations, compared to vehicle-exposed controls (Fig. 2B). When EVs derived from cisplatin-treated cells (control media, 0.5 µM or 10 µM cisplatin) were incubated with treatment-naïve cells, they demonstrated altered uptake by treatment-naïve cells. Specifically, EVs from GC1-spg cells exposed to 10 µM cisplatin showed a 5.7-fold increase in uptake by treatment-naïve GC1-spg cells (p=0.007) (Fig. 2C), while EVs from TM4 cells exhibited a 4.5-fold increase in uptake at 10 µM cisplatin, when incubated with treatment-naïve TM4 cells (p=0.02) (Fig. 2D). Representative images showing the uptake of fluorescently labelled EVs by TM4 treatment-naïve cells, incubated with EVs released from TM4 cells treated with different doses of cisplatin, are depicted in Fig. 2 (E-M). These conditions include control media (2E, F, G), 0.5 µM cisplatin (2H, I, J), and 10 µM cisplatin (K, L, M). Additionally, a single experiment tracking GC1-spg control EVs within a recipient GC1-spg cell at 24 hours post-incubation demonstrated that the EVs were distributed throughout the cell, with most moving at an intracellular speed of less than 0.05 µM/s (Supplementary Fig. 1C, 1D).

**Figure 2.**
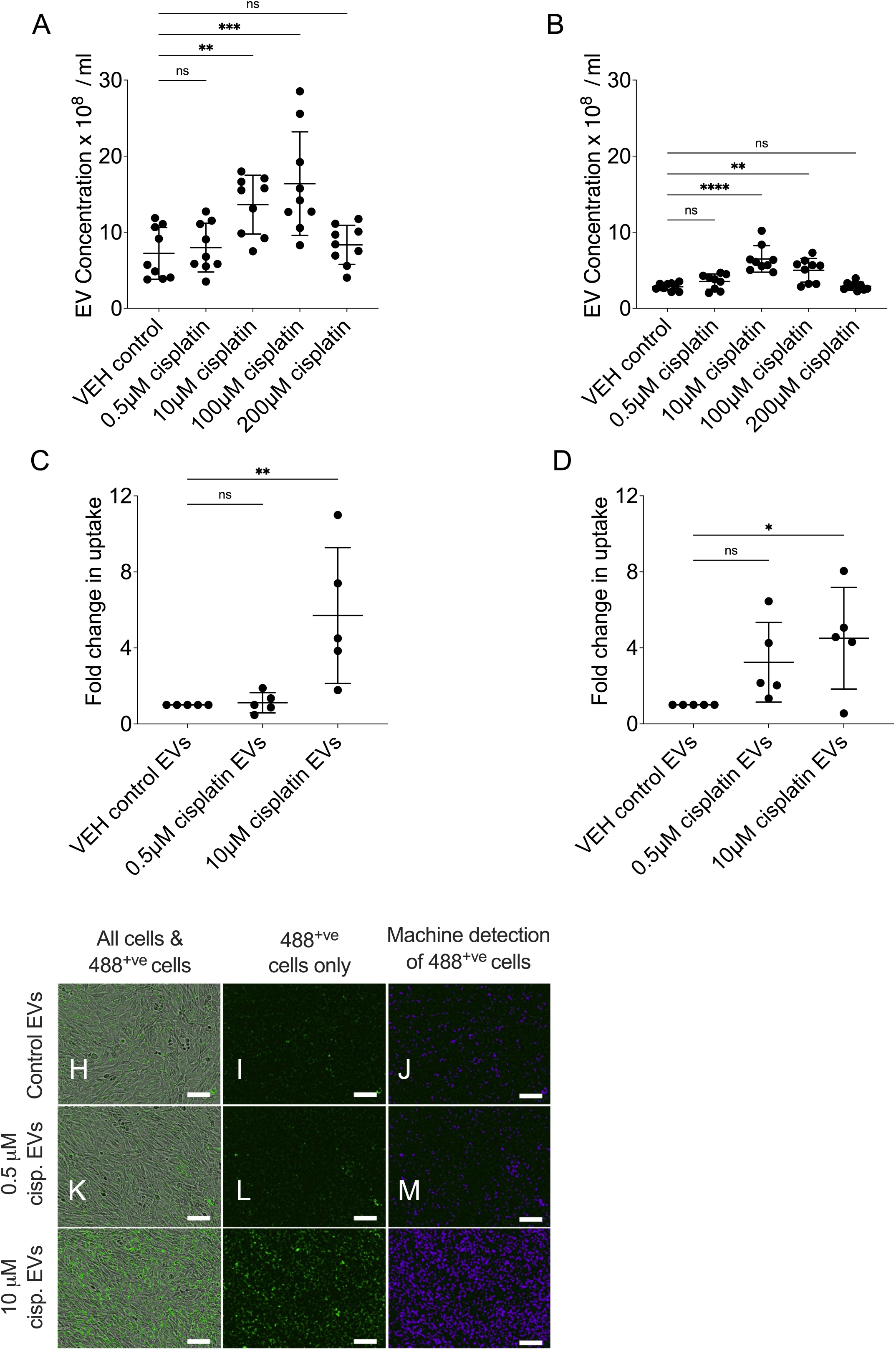
Quantification of EV release by cells treated with cisplatin versus VEH control in A) GC1-spg cells and B) TM4 cells. Quantification of cisplatin-induced EV uptake by C) GC1-spg cells and D) TM4 cells. Representative images of TM4 cells incubated with fluorescently labelled TM4 EVs released by cells treated with control media (E,F,G), 0.5 μΜ cisplatin (H,l,J), or 10 HM cisplatin (K,L,M). The first column of images shows cells with fluorescent EVs, the middle column shows fluorescent EVs only (green), and the right-hand column shows the same image but with the ‘mask’ used to detect fluorescent apoptosis (purple). Scale bar - 100 pm.

### 3.3 Cisplatin-derived TM4 EVs cause an increase in apoptosis in chemotherapy naïve TM4 cells

To investigate the apoptotic effects of EVs released by TM4 cells exposed to cisplatin at concentrations ranging from 0.5 μM - 200 μM, EVs were incubated with treatment-naïve TM4 cells for 24 h. A 2.3-fold increase in apoptosis was observed in cells incubated with EVs from cells exposed to 10 μM cisplatin, compared to those treated with control EVs (p=0.007) (Fig. 3A). However, no changes in apoptosis levels were detected in cells incubated with EVs from cells exposed to other cisplatin concentrations (0.5 μM, 100 μM, and 200 μM). Increasing the ratio (1x, 2.5x and 5x) of EVs released by TM4 cells treated with 10 μM cisplatin, then incubating them with treatment-naïve TM4 cells did not lead to a dose-dependent increase in apoptosis compared to controls, but rather maintained similar levels of apoptosis across experimental conditions (4.7-fold, 6.3-fold and 5.6-fold respectively, p<0.05) (Fig. 3B). Representative images of treatment naïve TM4 cells incubated with EVs released by TM4 cells treated with a range of cisplatin doses is presented in Fig. 3.

**Figure 3.**
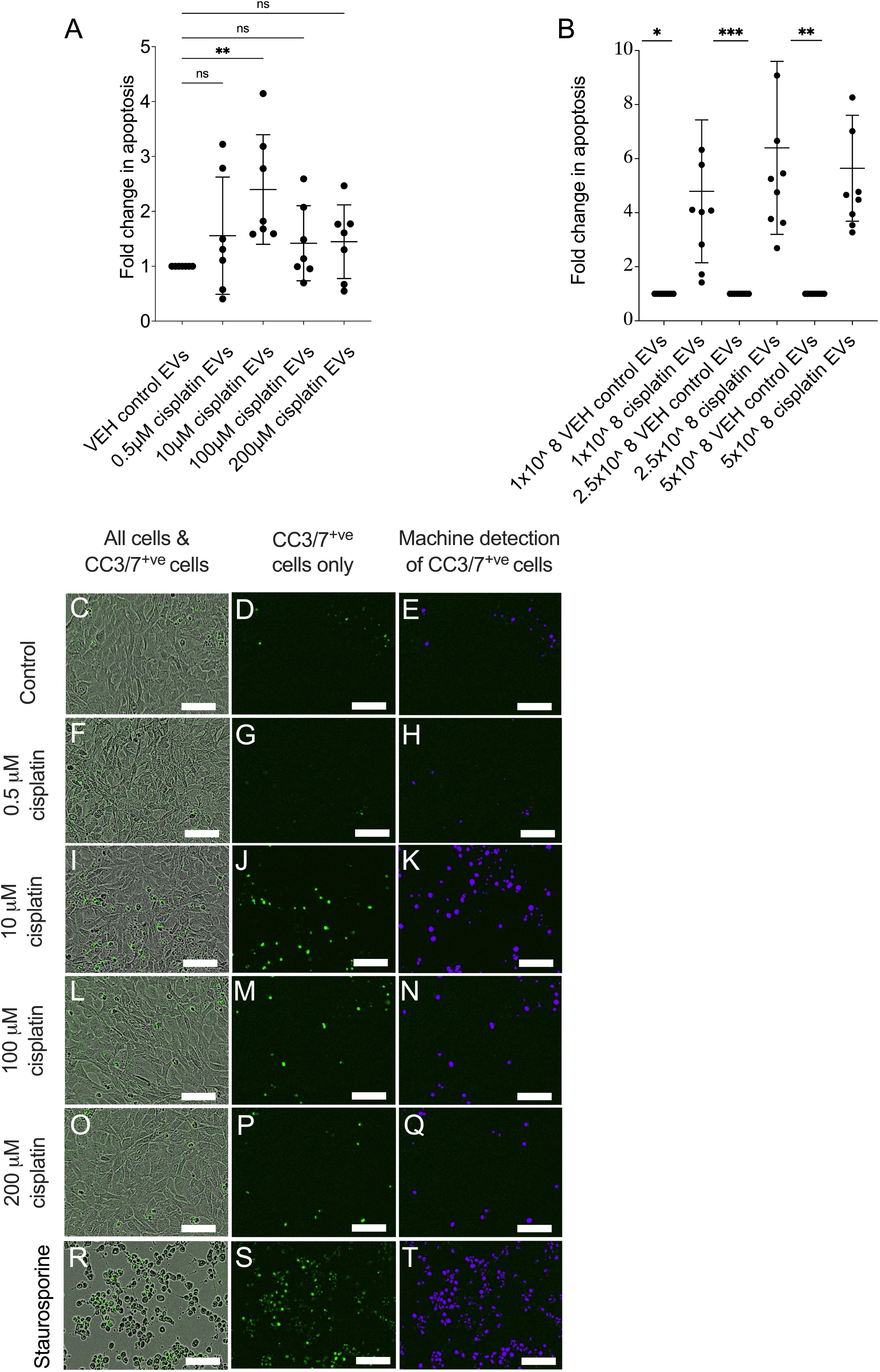
EV-mediated apoptosis in treatment-naïve TM4 cells incubated with EVs released by cisplatin-exposed TM4 cells. A) Quantification of apoptosis in treatment-naïve TM4 cells incubated with EVs released by cisplatin-treated TM4 cells (0-200µM cisplatin). B) Quantification of apoptosis in treatment-naïve TM4 cells incubated with an increasing ratio of EVs released by TM4 cells treated with 10µM cisplatin. * = p <0.05, ** p <0.01, ***p <0.005. Representative images of TM4 cells (grey) and fluorescent CC3/7 ^+ve^ apoptotic cells (green), fluorescent CC3/7 ^+v^θ apoptotic cells only (green) and machine learning detection of fluorescent CC3/7 ^+ve^ apoptotic cells (purple) in cells incubated with (C,D,E) EVs released by TM4 cells incubated with control media, (F,G,H) EVs released by TM4 cells treated with O.5µM cisplatin, (l,J,K) EVs released by TM4 cells treated with 10µM cisplatin, (L,M,N) EVs released by TM4 cells treated with 100µM cisplatin, (O,P,Q) EVs released by TM4 cells treated with 2OOµM cisplatin and (R,S,T) staurosporine-induced apoptosis positive control.

### 3.4 TM4-derived EVs reduce apoptosis in treatment-naïve GC1-spg cells and are chemoprotective to cisplatin

As previous data indicated that EVs released by TM4 cells exhibit a pro-apoptotic phenotype (Fig. 3A and 3B), we sought to investigate if this persisted when TM4 EVs were incubated with treatment-naïve GC1-spg cells. Using a range of cisplatin doses to generate TM4 EVs (0.5 µM, 10 µM, 100 µM and 200 µM), we observed a reduction in apoptosis in GC1-spg cells incubated with TM4 EVs (0.63-fold change, 0.63-fold change and 0.59-fold change for 0.5 µM, 10 µM, 100 µM doses of cisplatin, respectively, p<0.05), compared to those exposed to control EVs, except for EV released by TM4 cells treated with 200 μM cisplatin (0.73-fold change, p=0.15) (Fig. 4A). To investigate the potential chemoprotective effects of TM4 EVs to GC1-spg cells, we assessed whether these EVs incubated with GC1-spg cells, could mitigate cisplatin-induced apoptosis. To this end, we incubated EVs released by TM4 cells treated with control, 0.5 μM and 10 μM cisplatin with treatment-naïve GC1-spg cells for 24 h, followed by 10 μM cisplatin for a further 24 h. In all conditions, GC1-spg cells incubated with TM4 EVs showed lower levels of apoptosis compared to GC1-spg cells treated with cisplatin alone, specifically VEH control EVs (82.5 vs 39.2 apoptotic cells / mm^2^, p=0.004), 0.5 μM cisplatin EVs (82.5 vs 34.6 apoptotic cells / mm^2^, p=0.002) and 10 μm cisplatin EVs (82.5 vs 42.5 apoptotic cells / mm^2^, p=0.008) (Fig. 4B). These results indicate that TM4-derived EVs, whether released following exposure to cisplatin or in normal culture, confer chemoprotection to GC1-spg cells from cisplatin.

**Figure 4:**
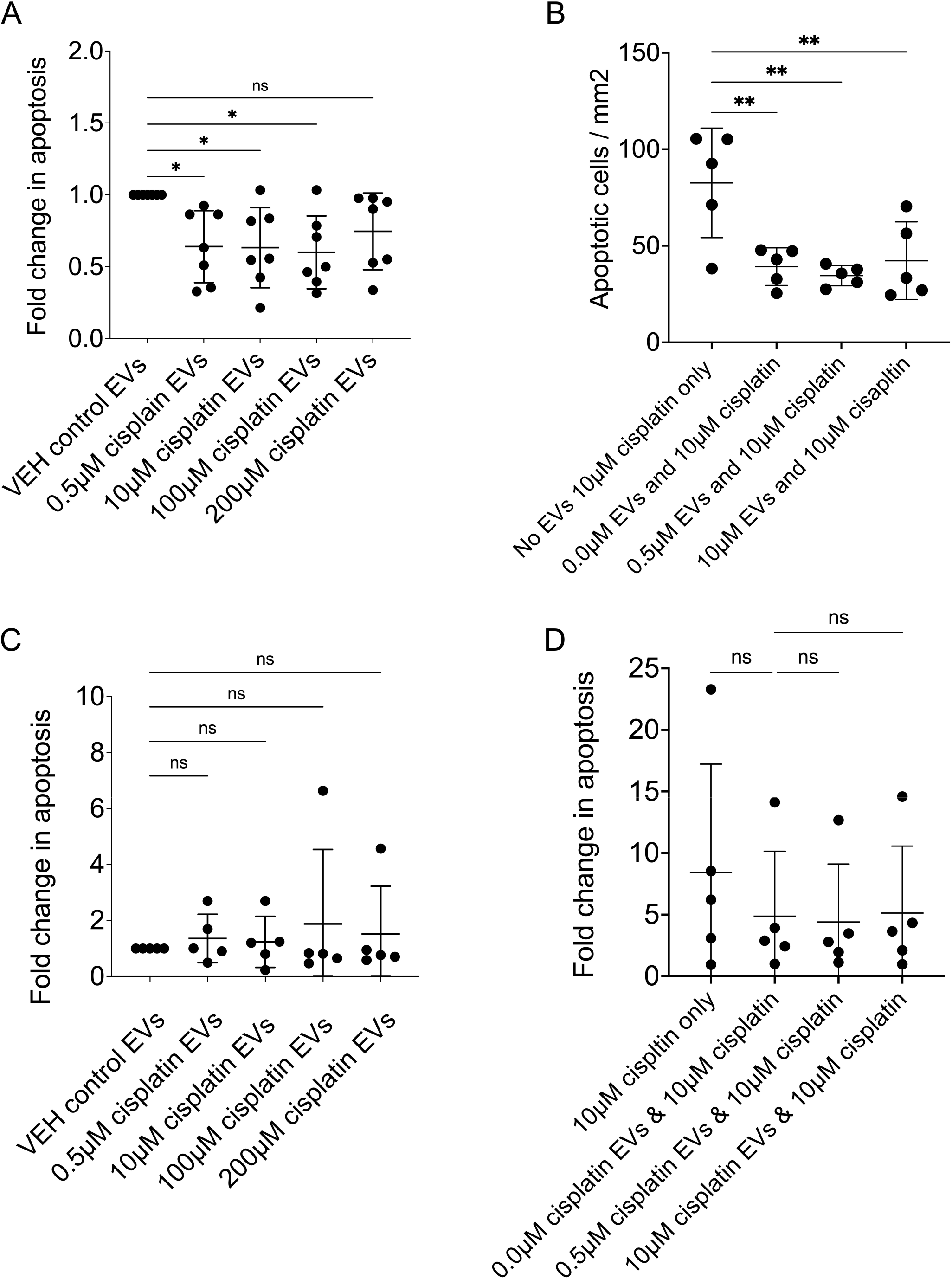
EV-mediated apoptosis and its reversal in TM4 and GC1-spg cells. A) Fold change in apoptosis in GC1-spg cells incubated with TM4 EVs released by cells cultured in various cisplatin doses, showing a decrease in apoptosis. B) Quantification of apoptotic cell number in GC1-spg cells incubated with TM4 EVs for 24 hours prior to a further 24 hours of cisplatin treatment, showing a reduction in apoptosis compared to control cells not incubated with EVs. C) Fold change in apoptosis in GC1-spg cells incubated with GC1-spg EVs released by cells treated with at a range of cisplatin doses, showing no difference compared to control EVs. D) Fold change in apoptosis in GC1-spg cells incubated with GC1-spg EVs for 24 hours prior to an additional 24 hours of cisplatin treatment, showing no change in apoptosis proportion compared to control cells not incubated with EVs. All data are presented as mean ± SD, * = p <0.05, ** = p <0.01.

### 3.5 GC1-spg EVs do not induce apoptosis in treatment naïve GC1-spg cells and are not chemoprotective

To investigate the impact of EVs released by GC1-spg cells exposed to various doses of cisplatin (0 - 200 µM) on treatment-naïve GC1-spg cells, GC1-spg EVs were incubated with treatment-naïve GC1-spg cells, and the levels of apoptosis were quantified over a 24-hour period. No change in apoptosis was observed in GC1-spg cells incubated with EVs released by cisplatin-exposed GC1-spg cells, compared to controls (Fig. 4C). To investigate the potential chemoprotective potential of these EVs, GC1-spg EVs released by cells treated with control, 0.5 μM and 10 μM cisplatin were incubated with GC1-spg cells for 24 hours, followed by 24 hours treatment with 10 μM of cisplatin and resulted in no fold change in apoptosis compared to the controls (Fig 4D).

### 3.6 Differential protein expression in TM4 cells exposed to cisplatin

Given the pro-apoptotic phenotype observed in EVs released by TM4 cells exposed to cisplatin, we sought to better understand the mechanisms underlying this phenomenon, by undertaking proteomics of EVs released by TM4 cells treated with 10 μM cisplatin or control media. Proteomics of TM4 EVs released by cells exposed to either cisplatin (10 μM) or control media demonstrated over 8,215 individual proteins mapping to 5527 genes with known functions (Fig. 5). These proteins included commonly accepted EV markers such as CD9, CD63, CD81 and CD82 as well as TSG-101 and heat shock protein-70, some of which were also identified either through Western Blot or single EV dSTORM imaging (Fig. 1E-J). Gene ontology analysis of these proteins identified 5,527 potential functions across 11 domains (Fig. 5A), with the largest domains being binding (40.8%) and catalytic activity (34.8%). Further analysis mapping the 8,215 individual proteins to biological processes identified 7121 genes with 12,489 potential functions. The largest classification of these potential functions included cellular processes (33.5%), metabolic processes (20.9%) and biological regulation (14.5%) (Fig. 5B)

**Figure 5:**
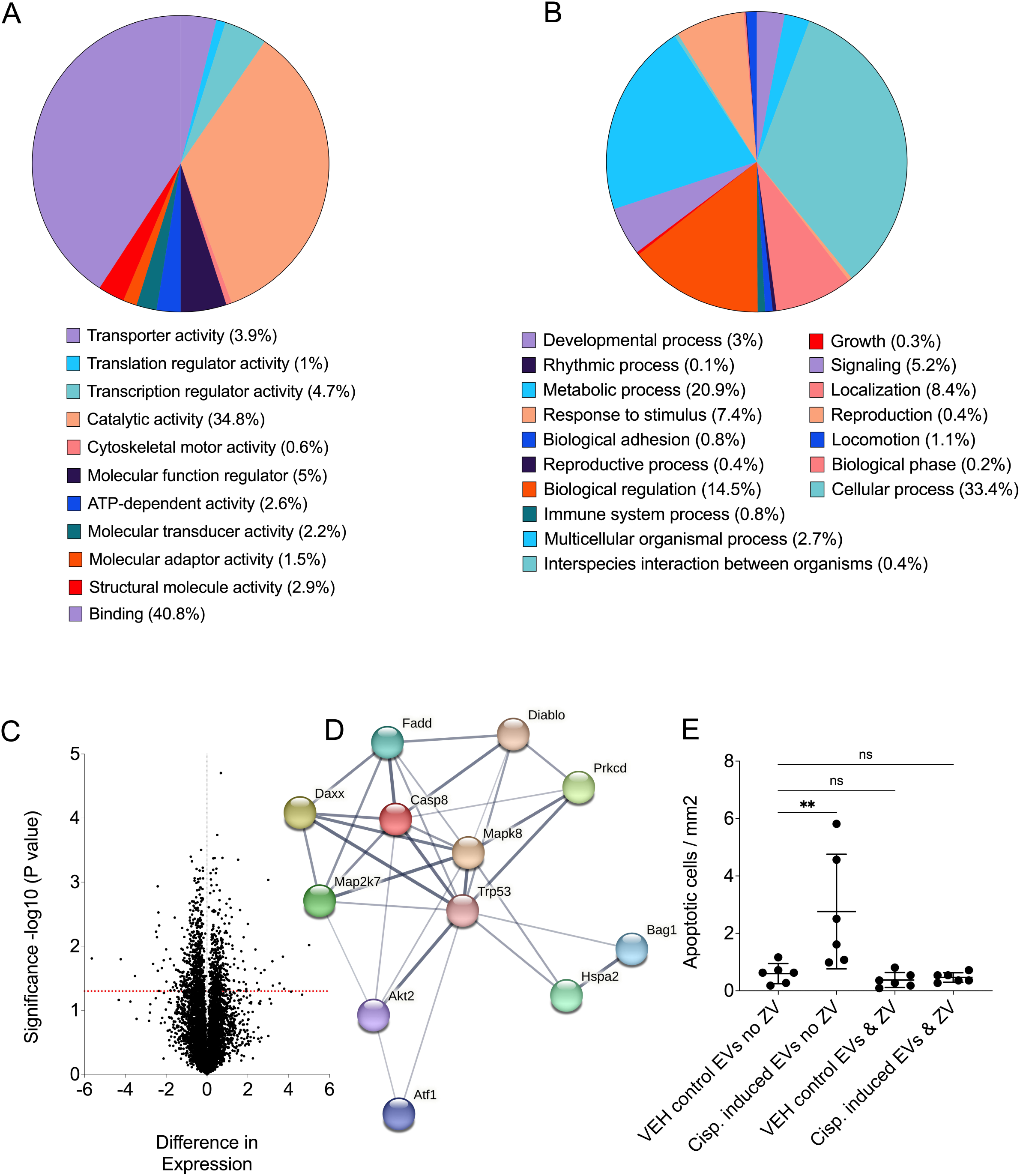
Proteomics of TM4 EV cargoes. A) Gene ontology of molecular functions identified within TM4 EVs. B) Gene ontology of biological processes within TM4 EVs. C) Volcano plot of TM4 EV cargoes comparing EVs released by cells exposed to control media versus cells exposed to 10µM cisplatin, with a Log 1.3 (p=0.05) indicated by the red dotted line. D) Search tool for the retrieval of interacting genes and proteins (STRING) of significantly differentially expressed proteins within TM4 EVs involved in apoptosis, showing multiple known interactions represented by lines connecting genes. E) Quantification of apoptotic cell number in treatment-naïve TM4 cells and the reversal of this with both control and cisplatin-induced EVs incubated with zVAD or PBS for 1 hour prior to incubation, demonstrating reversal of apoptosis.

Quantification of the expression of proteins within EVs released by cells exposed to cisplatin (10μM) or control media identified over 1,195 to be significantly differentially expressed (Fig. 5C). Interactions between these differentially expressed proteins were explored further using gene ontology analysis, which were mapped to 94 known distinct biological pathways (Supplementary Table 3). These included pathways such as Wnt signalling, gonadotrophin-releasing hormone receptor, integrin signalling and apoptosis signalling. Given that cisplatin mediates its cytotoxic effects through damaged DNA by forming DNA cross-links, we interrogated our proteomics data for proteins with a known role in DNA repair, identifying 36 proteins in total (Supplementary Table 4). These proteins accord with the chemoprotective effects observed in GC1-spg cells following delivery of Sertoli cell-derived cargoes.

### 3.7 Cisplatin exposure of TM4 cells leads to the production of EVs with upregulated pro-apoptotic proteins

Given the known gonadotoxic effects of chemotherapeutics such as cisplatin within the pre-pubertal testis (Matilionyte et al., 2022a, Tharmalingam et al., 2020, Smart et al., 2018), the potential effects on life-long fertility potential, and data pertaining to a pro-apoptotic phenotype in Fig. 3 and 4, we further explored the impact of cisplatin exposure on apoptosis signalling pathways within EVs released by TM4 cells.

Of the 8,215 proteins identified, 75 had a known function in apoptosis (Supplementary Table 5). Of these, 12 were differentially expressed, and all were upregulated within EVs which exhibited a pro-apoptotic phenotype when incubated with treatment-native TM4 cells (Fig 3A, 3B). STRING analysis of these proteins revealed a significant number of known interactions between these apoptotic proteins (Fig. 5D).

### 3.8 Reversal of EV-mediated apoptosis by caspase inhibition

Based on our finding that several caspases were identified as differentially expressed and upregulated within EVs released by chemotherapy-exposed cells based on our proteomics data, we hypothesised that caspase-mediated apoptosis may underpin the process.

To interrogate this mechanism, we used the pan-caspase inhibitor zVAD, incubated with EVs, to reverse the pro-apoptotic phenotype of EVs released by TM4 cells treated with cisplatin. TM4 cells were treated with either 10 µM cisplatin or control media to generate EVs over 24 hours, and these EVs were collected as previously described and incubated with zVAD. Following this, SEC was used to remove free zVAD and cell debris. EVs released from TM4 cells treated with cisplatin, then incubated with zVAD or control, were incubated with treatment-naïve TM4 cells for 24 hours. zVAD incubation with EVs inhibited their pro-apoptotic phenotype and EV-mediated apoptosis as compared to untreated EVs (Fig 5E).

Specifically, treatment-naïve TM4 cells incubated EVs without zVAD treatment demonstrated increased levels of apoptosis (0.59 vs 2.85 apoptotic cells / mm^2^, p=0.004) compared to zVAD-treated EVs (0.26 vs 0.46 apoptotic cells / mm^2^, p=0.9).

### 3.9 zVAD inhibits apoptotic cargoes within EVs and not recipient cells

To investigate if the reversal of apoptosis was mediated by unbound zVAD carried within EVs and delivered to recipient cells or by inhibition of apoptosis machinery within EVs, TM4 cells were incubated with zVAD-treated EVs for 24 hours, followed by treatment with cisplatin to induce apoptosis.

Cisplatin exposure significantly increased TM4 cell death at 24 hours, by over two-fold, compared to control, p<0.05 (Supplementary Figure 2). Pre-incubation with zVAD-treated EVs did not protect TM4 cells from subsequent cisplatin-induced apoptosis for both cell incubated with VEH control EVs (10.73 vs. 7.73 apoptotic cells / mm^2^, p>0.39), or 10 μM cisplatin EVs (10.73 vs. 6.99 apoptotic cells / mm^2^, p>0.25). These findings show that although EV incubation with zVAD reverses the pro-apoptotic phenotype of the EVs, potentially by inhibiting their pro-apoptotic cargoes, the inherent caspase-mediated apoptotic machinery within the cells remains unchanged. These data would imply the apoptotic impact of these EVs is mediated by the caspase machinery within the EVs delivered to treatment-naïve Sertoli cells and that EVs do not carry unbound zVAD within them.

## 4 Discussion

We investigated the role of EV-mediated intercellular communication between germ and somatic cells in the prepubertal testis using model cell lines. We used the chemotherapeutic agent cisplatin to recapitulate chemotherapy-induced gonadotoxicity in the testis, as occurs in childhood cancer treatment. This study aimed to explore potential mechanisms through which EVs can modulate post-treatment damage within the testis.

Our findings demonstrate increased EV release from germ and Sertoli cells following chemotherapy, with alterations in recipient cell interactions and EV impact. These changes likely reflect altered cell homeostasis and subsequent EV-mediated cell-to-cell communication. EVs from chemotherapy-treated Sertoli cells contain pro-apoptotic cargoes, increasing apoptosis in treatment-naïve Sertoli cells. Conversely, TM4 cell-derived EVs protect germ cells from cisplatin-induced apoptosis. Both EV cargo and uptake were altered, with a four-fold increase in uptake of EVs from cisplatin-treated TM4 cells, indicating potential changes in EV surface molecules impacting their interaction and uptake by recipient cells.

Considerable attention has been paid to direct chemotherapy-induced damage to germ and Sertoli cells in the testis. However, the present data provide evidence for secondary effects of chemotherapy, mediated by Sertoli cell-derived EVs which can impact on neighbouring germ and Sertoli cells. Our data indicate that Sertoli-Sertoli cell communication via EVs can propagate chemotherapy-induced testicular damage, whilst Sertoli cell-derived EVs can act as chemoprotectants, reducing levels of germ cell apoptosis.

### 4.1 Sertoli-Sertoli cell communication & mechanisms of EV-mediated gonadotoxicitiy

Sertoli cells provide a specialised immune-privileged microenvironment within the seminiferous tubules of the testis, which supports germ cell differentiation from SSC to mature sperm (Qu et al., 2020). Sertoli cells regulate the blood-testis barrier, protecting developing germ cells from deleterious molecules which may damage and perturb germ cell development. Activated by follicle stimulating hormone, they release hormones and other signalling moieties, including EVs, that promote germ cell growth and differentiation through regulation of the microenvironment (Gerber et al., 2016, Wang et al., 2022).

Exposure to cisplatin resulted in approximately 10% of Sertoli cells undergoing apoptosis. Thus, the EVs carrying pro-apoptotic machinery, capable of inducing apoptosis in treatment-naïve Sertoli cells, were derived from either those undergoing apoptosis and propagating this effect through the action of the EVs they released or from the surviving, but potentially damaged, Sertoli cells. This horizontal transfer of cargo, particularly a network of pro-apoptotic machinery including caspases, appears to trigger a cascade effect, amplifying the initial chemotherapeutic insult and contributing to widespread Sertoli cell loss. Furthermore, compared to control EVs, cisplatin-induced Sertoli cell EVs were taken up at higher levels (four times) compared to control EVs, further amplifying the delivery of pro-apoptotic cargoes.

Caspase 8 was identified as significantly elevated within our EVs along with its trigger, Fas cell surface death receptor, a member of the TNF receptor superfamily (Fu et al., 2016). While caspase 3 and 7 were not upregulated in EVs after chemotherapy exposure, we hypothesise that the relative increase in cisplatin-derived EV uptake results in a significant increase in the payload of pro-apoptotic machinery delivered to recipient cells, resulting in increased apoptosis.

Investigating the mechanism of apoptosis, we hypothesised this may be mediated by caspases, given these were upregulated in EVs and the primary readout of apoptosis used in these experiments. We used the pan-caspase inhibitor zVAD to inhibit caspase-based cargoes within EVs and were able to reverse their pro-apoptotic phenotype when incubated with recipient cells. We further explored the potential for zVAD to be transferred to cells within EVs via a Trojan horse mechanism; however, incubation of TM4 cells with zVAD-treated EVs did not prevent them from undergoing apoptosis in response to chemotherapy. These data suggest the apoptosis mediated by EVs is due to caspase-based cargoes within the EVs.

### 4.2 Sertoli-germ cell communication & the chemoprotective effect of Sertoli cell EVs on germ cells

The present study indicates that communication between Sertoli cells and germ cells via EVs plays a crucial protective role for germ cells, including during recovery after chemotherapy exposure. In the adult testis, the role of Sertoli cells in germ cell survival is well established. They secrete numerous factors, including EV-based, which promote testis function as well as germ cell development and survival (Li et al., 2021, Chen et al., 2023, O’Donnell et al., 2022, Wang et al., 2023). As with other body systems, cargo loading into EVs in the testis is dynamic and under hormonal regulation (Mancuso et al., 2018) with changes observed in response to hypoxia (Ma et al., 2021) and infection (He et al., 2019).

While some studies have reported the protective roles of EVs in the adult testis, there is limited data regarding their function in the prepubertal testis. The few studies that have characterised pre-pubertal Sertoli cell EVs in the human testis have identified several RNAs and proteins targeting genes that regulate spermatogenesis, the cell cycle, and immune regulation (Tan et al., 2022). Their crucial role in germ cell survival has been reported in porcine models, where germ cell proliferation was impaired in response to inhibition of EV-uptake (Thiageswaran et al., 2022). Importantly, these studies did not involve EVs derived from chemotherapy-exposed cells.

The mechanisms by which these EVs mediate protection to germ cells have not been explored; however, of the 1195 significantly differentially expressed proteins within Sertoli cells treated with cisplatin, 36 with known roles in DNA repair were identified. Cisplatin mediates DNA damage through the formation of DNA crosslinks. Therefore, it is possible that Sertoli cell EV cargoes carry repair messages that augment their innate repair mechanisms resulting in the novel chemo-protective effects identified in the present study.

These proteins include DNA repair protein RAD51 homolog 3 (RAD51), which plays a central role in facilitating accurate repair of double-strand breaks. Enhancing homologous recombination efficiency may be achieved through EV-mediated delivery of additional RAD51 and its key mediators, including the replication factor C subunit 5 (RFC1-5) subunits, promoting high-fidelity repair in germ cells and preventing the accumulation of cytotoxic double-strand breaks (Bhattacharya et al., 2017). Another EV-mediated protein conferring protection to germ cells is nucleotide excision repair, which is primarily responsible for removing bulky DNA adducts caused by cisplatin (Friedberg, 2001). In addition to DNA repair, nucleotide excision repair has been implicated in the development of cisplatin resistance in ovarian cancer, demonstrating its potential protective effect to germ cells prior to cisplatin exposure (Ferry et al., 2000).

DNA repair proteins significantly upregulated in EVs released by cisplatin-treated Sertoli cells include RAD51 homolog 3 (RAD51), which plays a central role in facilitating accurate repair of double-strand breaks. Enhancing homologous recombination efficiency may be achieved through EV-mediated delivery of additional RAD51 and its key mediators, including the replication factor C subunit 5 (RFC1-5) subunits. This can promote high-fidelity repair in germ cells and prevent the accumulation of cytotoxic double-strand breaks (Bhattacharya et al., 2017). Another EV-mediated protein upregulated in cisplatin-treated Sertoli cell EVs, possibly conferring protection to germ cells, is nucleotide excision repair, which is primarily responsible for removing bulky DNA adducts caused by cisplatin (Friedberg, 2001). In addition to DNA repair, nucleotide excision repair has been implicated in the development of cisplatin resistance in ovarian cancer, demonstrating its potential protective effect to germ cells prior to cisplatin exposure (Ferry et al., 2000).

Poly (ADP-ribose) polymerase 2 (PARP2), which promotes DNA repair, was also upregulated in cisplatin-derived TM4 EVs. PARP2 is activated in response to DNA strand breaks and recruits additional repair factors, safeguarding genomic DNA in response to cisplatin-mediated damage (Amé et al., 1999). Another protein delivered by Sertoli cell EVs is Msh3, a DNA mismatch repair protein that recognises and binds to bulky DNA lesions resulting from cisplatin damage. While Msh3 does not directly excise damaged DNA, its binding to interstrand cross-links is vital for initiating subsequent repair processes. These include nucleotide excision repair and homologous recombination, both of which are crucial for resolving cross-links and repairing double-strand breaks as outlined above. Moreover, Msh3 deficiency increases cellular sensitivity to cisplatin, which leads to poor repair of double-strand breaks and the accumulation of toxic DNA lesions, ultimately resulting in apoptosis (Sawant et al., 2015).

These data add to the emerging body of evidence suggesting that Sertoli cell-derived EVs promote both germ cell development and act as potential chemoprotectants (Ma et al., 2023). Through the targeted delivery of protective cargos, these EVs may function as natural chemoprotectants, giving us exciting insights into endogenous pathways within the testis that attempt to reduce the damage caused by chemotherapy. Fully elucidating the pathways which are implicated in this chemoprotection is key to further manipulation to better protect the testis from unwanted, off-target effects of chemotherapy-mediated gonadotoxicity in childhood cancer.

### 4.3 Limitations

While much of this work focuses on the impact of EVs, it is crucial to consider the activation state of the recipient cell, which can significantly influence EV interaction, uptake, and subsequent fate. The cell’s activation status can determine whether internalised EVs are directed towards degradation in the lysosome, recycled via multivesicular bodies, or ultimately release their cargo to modulate cell function. Understanding both the impact of EVs and the activation state of recipient cells is needed to fully elucidate the mechanisms through which they work. Our data give us valuable insights into the complex environment of the prepubertal testis and the role of EV-mediated communication, paving the way for further work and even potential therapeutic interventions in the field of fertility preservation for childhood cancer. The approach used in the present study allowed for the separation of the direct effects of chemotherapy on testicular cells from indirect secondary effects mediated by EVs. However, this required the use of immortalised juvenile mouse cell lines, including TM4 cells (Sertoli) and GC1-spg (Type B spermatogonia). Further validation using human testicular cells would provide further evidence for the clinical relevance of these findings.

While our in vitro system may not fully replicate the complex environment of the testis, it provides a platform which we can manipulate to characterise the effects of Sertoli cell EVs on recipient Sertoli and germ cells. This reductionist approach is needed in order to investigate individual steps in EV release and their impact on recipient cells. Focusing on cisplatin, a widely used chemotherapeutic agent, offers insights into the consequences of chemotherapy on testicular function. Although the response to other chemotherapeutics or EVs derived from different exposures remains unknown, this approach advances our understanding of the mechanisms underlying EV-mediated communication in the testis. These findings contribute to a foundational understanding that could inform strategies to mitigate chemotherapy-induced reproductive toxicity.

## 5 Conclusion

We have identified a dual role of Sertoli cell-derived EVs in the testis, which is dependent on the recipient cell population. Our data demonstrate that cisplatin-induced EVs from Sertoli cells can induce apoptosis among Sertoli cells, while simultaneously conferring chemoprotection to germ cells. Given that the role of Sertoli cells is to protect developing cells, their EVs also convey protective effects to germ cells, which may be an extension of this function. The pro-apoptotic effect on Sertoli cells, mediated through caspase-dependent pathways, represents a previously unrecognized mechanism of secondary testicular damage following chemotherapy. This “second hit” phenomenon suggests that initial chemotherapy exposure triggers a cascade of EV-mediated damage that extends beyond direct cytotoxicity. Conversely, the chemoprotective effect on germ cells, potentially mediated through DNA repair proteins including RAD51, RFC1-5, PARP2, and Msh3, suggests an endogenous protective mechanism that could be harnessed therapeutically.

Better understanding and characterisation of the full impact of cisplatin in the prepubertal testes is essential for manipulating these pathways and ultimately reducing the deleterious effects of chemotherapy. This knowledge could lead to novel interventions to preserve the future fertility of children with cancer. Recognising that Sertoli cell-derived EVs may naturally attempt to protect germ cells from chemotherapy damage opens the possibility to manipulate these endogenous effects of EVs to deliver protective factors to the testis prior to chemotherapy treatment. This needs to be balanced against detrimental impacts to Sertoli cells, however. The identification of caspase-mediated apoptosis as a mechanism for protection provides a potential target for developing therapeutics to minimise gonadotoxicity. This work establishes EVs as important mediators of testicular response to chemotherapy and demonstrates novel insights into their mechanistic impact within the prepubertal testes, which may provide therapeutic targets for preserving fertility in childhood cancer patients.

## Acknowledgements

Electron microscopy was performed in the Microscopy Core of the Program in Membrane Biology by Diane Capen who we would like to thank for her help with this work.

The authors would like to thanks the following for their assistance with this work: David Greenald, Federica Lopes, Gabriele Matilionyte, Christos Spanos, Sara Veiga, Evelyn Luciani, Daniel Ruiz Torres and Uyen Ho.

## Data Availability Statement

Data is available on request from the corresponding author.

