## Supplementary Tables for "Chemotherapy-induced transfer of apoptotic machinery in extracellular vesicles between somatic and germ cells of the testis: mechanistic insights into onco-fertility preservation in pre-pubertal boys"

**Supplementary Table 1** - Primary and secondary antibodies used for Western blotting.

| Antibody | Primary / Secondary | Cat No. | Manufacturer | Dilution | Expected Weight |
| --- | --- | --- | --- | --- | --- |
| Recombinant Rabbit anti-TSG-101 | Primary | AB125011 | Abcam | 1 in 5000 | 45 kDa |
| Rat anti-tubulin | Primary | AB6160 | Abcam | 1 in 5000 | 55 kDa |
| IRdye 800CW Goat anti-Rabbit IgG  (secondary for TSG-101) | Secondary | 926-32211 | LiCor | 1 in 15,000 | NA |
| IRdye 680CW Goat  anti-rat IgG  (secondary for alpha tubulin) | Secondary | 926-68076 | LiCor | 1 in 15,000 | NA |

**Supplementary Table** **2** - Inclusion mass list and corresponding isolation windows for mass spectrometry

| m/z | z | t start (min) | | t stop (min) | | Isolation Window (m/z) | |
| --- | --- | --- | --- | --- | --- | --- | --- |
| m/z | z | | t start (min) | | t stop (min) | | Isolation Window (m/z) |

| 383.375 | 3 | 0 | 185 | 66.8 |
| --- | --- | --- | --- | --- |
| 423 | 3 | 0 | 185 | 13.5 |
| 435 | 3 | 0 | 185 | 11.5 |
| 446.5 | 3 | 0 | 185 | 12.5 |
| 458 | 3 | 0 | 185 | 11.5 |
| 469 | 3 | 0 | 185 | 11.5 |
| 480 | 3 | 0 | 185 | 10.5 |
| 490.5 | 3 | 0 | 185 | 10.5 |
| 501 | 3 | 0 | 185 | 11.5 |
| 512 | 3 | 0 | 185 | 11.5 |
| 523 | 3 | 0 | 185 | 11.5 |
| 533.5 | 3 | 0 | 185 | 10.5 |
| 544 | 3 | 0 | 185 | 11.5 |
| 554.5 | 3 | 0 | 185 | 10.5 |
| 565 | 3 | 0 | 185 | 11.5 |
| 575.5 | 3 | 0 | 185 | 10.5 |
| 586 | 3 | 0 | 185 | 11.5 |
| 597.5 | 3 | 0 | 185 | 12.5 |
| 609.5 | 3 | 0 | 185 | 12.5 |
| 621.5 | 3 | 0 | 185 | 12.5 |
| 633 | 3 | 0 | 185 | 12.5 |
| 645 | 3 | 0 | 185 | 13.5 |
| 657.5 | 3 | 0 | 185 | 12.5 |
| 670.5 | 3 | 0 | 185 | 14.5 |
| 684 | 3 | 0 | 185 | 13.5 |
| 697 | 3 | 0 | 185 | 13.5 |
| 710.5 | 3 | 0 | 185 | 14.5 |
| 725.5 | 3 | 0 | 185 | 16.5 |
| 741 | 3 | 0 | 185 | 15.5 |
| 756.5 | 3 | 0 | 185 | 16.5 |
| 773.5 | 3 | 0 | 185 | 18.5 |
| 791 | 3 | 0 | 185 | 17.5 |
| 808.5 | 3 | 0 | 185 | 18.5 |
| 827 | 3 | 0 | 185 | 19.5 |
| 846.5 | 3 | 0 | 185 | 20.5 |
| 866.5 | 3 | 0 | 185 | 20.5 |
| 887.5 | 3 | 0 | 185 | 22.5 |
| 910.5 | 3 | 0 | 185 | 24.5 |
| 935.5 | 3 | 0 | 185 | 26.5 |
| 962.5 | 3 | 0 | 185 | 28.5 |
| 992 | 3 | 0 | 185 | 31.5 |
| 1025 | 3 | 0 | 185 | 35.5 |
| 1063 | 3 | 0 | 185 | 41.5 |
| 1108.5 | 3 | 0 | 185 | 50.5 |
| 1391.62 | 3 | 0 | 185 | 516.8 |

**Supplementary Table 3** - Biological processes identified within TM4 EVs.

| **Biological Process** | **Representation of total biological processes** |
| --- | --- |
| Wnt signaling pathway | 4.70% |
| Huntington disease | 3.90% |
| CCKR signaling map | 3.70% |
| Gonadotropin-releasing hormone receptor pathway | 3.40% |
| Integrin signalling pathway | 3.40% |
| Inflammation mediated by chemokine and cytokine signaling pathway | 3.40% |
| Parkinson disease | 3.20% |
| PDGF signaling pathway | 3.20% |
| EGF receptor signaling pathway | 3.20% |
| Apoptosis signaling pathway | 2.90% |
| Angiogenesis | 2.90% |
| FGF signaling pathway | 2.20% |
| Cadherin signaling pathway | 2.20% |
| Ubiquitin proteasome pathway | 2.00% |
| Alzheimer disease-presenilin pathway | 1.70% |
| p53 pathway | 1.70% |
| Transcription regulation by bZIP transcription factor | 1.70% |
| TGF-beta signaling pathway | 1.70% |
| Ras Pathway | 1.70% |
| General transcription regulation | 1.70% |
| Endothelin signaling pathway | 1.70% |
| Alzheimer disease-amyloid secretase pathway | 1.50% |
| T cell activation | 1.50% |
| FAS signaling pathway | 1.50% |
| DNA replication | 1.50% |
| Cell cycle | 1.50% |
| Axon guidance mediated by netrin | 1.20% |
| Cytoskeletal regulation by Rho GTPase | 1.20% |
| B cell activation | 1.20% |
| Axon guidance mediated by semaphorins | 1.00% |
| TCA cycle | 1.00% |
| PI3 kinase pathway | 1.00% |
| Muscarinic acetylcholine receptor 1 and 3 signaling pathway | 1.00% |
| Interleukin signaling pathway | 1.00% |
| p53 pathway feedback loops 2 | 1.00% |
| Heterotrimeric G-protein signaling pathway-Gq alpha and Go alpha mediated pathway | 1.00% |
| Hedgehog signaling pathway | 1.00% |
| Glycolysis | 1.00% |
| Histamine H1 receptor mediated signaling pathway | 1.00% |
| VEGF signaling pathway | 0.70% |
| Toll receptor signaling pathway | 0.70% |
| Nicotinic acetylcholine receptor signaling pathway | 0.70% |
| Muscarinic acetylcholine receptor 2 and 4 signaling pathway | 0.70% |
| Metabotropic glutamate receptor group I pathway | 0.70% |
| p53 pathway by glucose deprivation | 0.70% |
| Thyrotropin-releasing hormone receptor signaling pathway | 0.70% |
| Oxytocin receptor mediated signaling pathway | 0.70% |
| p38 MAPK pathway | 0.70% |
| Nicotine pharmacodynamics pathway | 0.70% |
| Dopamine receptor mediated signaling pathway | 0.70% |
| Pyruvate metabolism | 0.70% |
| 5HT2 type receptor mediated signaling pathway | 0.70% |
| Axon guidance mediated by Slit/Robo | 0.50% |
| Alpha adrenergic receptor signaling pathway | 0.50% |
| Adrenaline and noradrenaline biosynthesis | 0.50% |
| Leucine biosynthesis | 0.50% |
| De novo purine biosynthesis | 0.50% |
| Metabotropic glutamate receptor group II pathway | 0.50% |
| Metabotropic glutamate receptor group III pathway | 0.50% |
| ATP synthesis | 0.50% |
| Vasopressin synthesis | 0.50% |
| Heterotrimeric G-protein signaling pathway-Gi alpha and Gs alpha mediated pathway | 0.50% |
| Pyrimidine Metabolism | 0.50% |
| Blood coagulation | 0.50% |
| Toll pathway-drosophila | 0.20% |
| Ornithine degradation | 0.20% |
| Methylmalonyl pathway | 0.20% |
| Mannose metabolism | 0.20% |
| Isoleucine biosynthesis | 0.20% |
| Flavin biosynthesis | 0.20% |
| De novo pyrimidine deoxyribonucleotide biosynthesis | 0.20% |
| Oxidative stress response | 0.20% |
| Notch signaling pathway | 0.20% |
| Synaptic vesicle trafficking | 0.20% |
| GABA-B receptor II signaling | 0.20% |
| JAK/STAT signaling pathway | 0.20% |
| Alanine biosynthesis | 0.20% |
| Interferon-gamma signaling pathway | 0.20% |
| Adenine and hypoxanthine salvage pathway | 0.20% |
| Insulin/IGF pathway-protein kinase B signaling cascade | 0.20% |
| Insulin/IGF pathway-mitogen activated protein kinase kinase/MAP kinase cascade | 0.20% |
| Valine biosynthesis | 0.20% |
| Hypoxia response via HIF activation | 0.20% |
| P53 pathway feedback loops 1 | 0.20% |
| Heterotrimeric G-protein signaling pathway-rod outer segment phototransduction | 0.20% |
| Sulfate assimilation | 0.20% |
| General transcription by RNA polymerase I | 0.20% |
| Enkephalin release | 0.20% |
| Angiotensin II-stimulated signaling through G proteins and beta-arrestin | 0.20% |
| Histamine H2 receptor mediated signaling pathway | 0.20% |
| Beta2 adrenergic receptor signaling pathway | 0.20% |
| Beta1 adrenergic receptor signaling pathway | 0.20% |
| 5HT1 type receptor mediated signaling pathway | 0.20% |
| 5-Hydroxytryptamine degredation | 0.20% |

**Supplementary Table 4** - Differentially expressed proteins with a role in DNA repair identified in TM4 EVs.

| **Protein** | |
| --- | --- |
| Actin | Peptidylprolyl isomerase E |
| Actin-like protein 6A | Serine/threonine-protein phosphatase 5 |
| Cyclin-H | E3 ubiquitin-protein ligase RAD18 |
| Serine/threonine-protein kinase Chk2 | DNA repair protein RAD51 homolog 3 |
| TFIIH basal transcription factor complex helicase XPB subunit | Replication factor C subunit 1 |
| DNA repair endonuclease XPF | Replication factor C subunit 2 |
| DNA excision repair protein ERCC-5 | Replication factor C subunit 4 |
| Fanconi anemia group I protein | Replication factor C subunit 5 |
| E3 ubiquitin-protein ligase FANCL | Replication timing regulatory factor 1 |
| Alpha-ketoglutarate-dependent dioxygenase FTO | Replication protein A 70 kDa subunit |
| COP9 signalosome complex subunit 1 | Tyrosyl-DNA phosphodiesterase 1 |
| General transcription factor IIH subunit 4 | DNA topoisomerase 2-binding protein 1 |
| Histone acetyltransferase KAT5 | E3 ubiquitin-protein ligase TRIM25 |
| DNA ligase 1 | Tumor suppressor p53 |
| DNA mismatch repair protein | Tumor suppressor p53-binding protein 1 |
| Nuclear receptor SET domain-containing protein 2 | RNA polymerase II-associated factor 1 homolog |
| Poly(ADP-ribose) polymerase 2 | DNA repair protein complementing XP-C cells |
| DNA-directed RNA polymerases I, II, and III subunit RPABC3 | Transcriptional activator YY1 |

**Supplementary Table 5** - Proteins with a role in apoptosis identified in TM4 EVs.

| **Protein** | |
| --- | --- |
| Kelch-like protein 20 | Caspase-3 |
| Transcription factor p65 | Heat shock 70 kDa protein 1-like |
| Bcl-2-like protein 1 | Mitogen-activated protein kinase 1 |
| Mitogen-activated protein kinase kinase kinase kinase 2 | RAC-gamma serine/threonine-protein kinase |
| Cytochrome c, somatic | Transcription factor AP-1 |
| Mitogen-activated protein kinase kinase kinase kinase 4 | Death domain-containing protein CRADD |
| Tumor necrosis factor receptor superfamily member 1A | Apoptotic protease-activating factor 1 |
| Mitogen-activated protein kinase kinase kinase kinase 3 | Bcl-2-like protein 11 |
| Protein kinase C epsilon type | DNA replication licensing factor MCM5 |
| FAS-associated death domain protein | Heat shock-related 70 kDa protein 2 |
| Eukaryotic translation initiation factor 2 subunit 1 | Tumor necrosis factor receptor superfamily member 6 |
| TBC1 domain family member 13 | Diablo homolog, mitochondrial |
| Bcl-2-related ovarian killer protein | Mitogen-activated protein kinase 9 |
| Death domain-associated protein 6 | Phosphatidylinositol 4,5-bisphosphate 3-kinase catalytic subunit beta isoform |
| Cation-independent mannose-6-phosphate receptor | Tumor necrosis factor receptor superfamily member 10B |
| Cyclic AMP-dependent transcription factor ATF-6 beta | Apoptosis regulator Bcl-2 |
| Mitogen-activated protein kinase 3 | Interferon-inducible double-stranded RNA-dependent protein kinase activator A |
| NF-kappa-B inhibitor alpha | Apoptosis-inducing factor 1, mitochondrial |
| Caspase-7 | Cyclic AMP-dependent transcription factor ATF-1 |
| Mitogen-activated protein kinase kinase kinase kinase 5 | Caspase-8 |
| Tumor necrosis factor receptor superfamily member 23 | Proto-oncogene c-Rel |
| Mitogen-activated protein kinase 15 | TNF receptor-associated factor 6 |
| Transcription factor RelB | Mitogen-activated protein kinase 8 |
| Dual specificity mitogen-activated protein kinase kinase 7 | Phosphatidylinositol 4,5-bisphosphate 3-kinase catalytic subunit delta isoform |
| Caspase-9 | RAC-alpha serine/threonine-protein kinase |
| E3 ubiquitin-protein ligase XIAP | Inhibitor of nuclear factor kappa-B kinase subunit alpha |
| BAG family molecular chaperone regulator 4 | TNF receptor-associated factor 2 |
| Interferon-induced, double-stranded RNA-activated protein kinase | MAP kinase-activating death domain protein |
| BAG family molecular chaperone regulator 3 | RAC-beta serine/threonine-protein kinase |
| Bcl-2 homologous antagonist/killer | Nuclear factor NF-kappa-B p100 subunit |
| Receptor-interacting serine/threonine-protein kinase 1 | Mitogen-activated protein kinase kinase kinase 1 |
| Phosphatidylinositol 4,5-bisphosphate 3-kinase catalytic subunit alpha isoform | Dual specificity mitogen-activated protein kinase kinase 4 |
| Protein kinase C delta type | Nuclear factor NF-kappa-B p105 subunit |
| Tumor necrosis factor receptor type 1-associated DEATH domain protein | Bax inhibitor 1 |
| BAG family molecular chaperone regulator 1 | Inhibitor of nuclear factor kappa-B kinase subunit beta |
| Heat shock 70 kDa protein 13 | Heat shock cognate 71 kDa protein |
| Dual specificity mitogen-activated protein kinase kinase 3 | Apoptosis regulator BAX |
