## Supplementary figures and images for "Chemotherapy-induced transfer of apoptotic machinery in extracellular vesicles between somatic and germ cells of the testis: mechanistic insights into onco-fertility preservation in pre-pubertal boys"

### Supplementary Figure 1

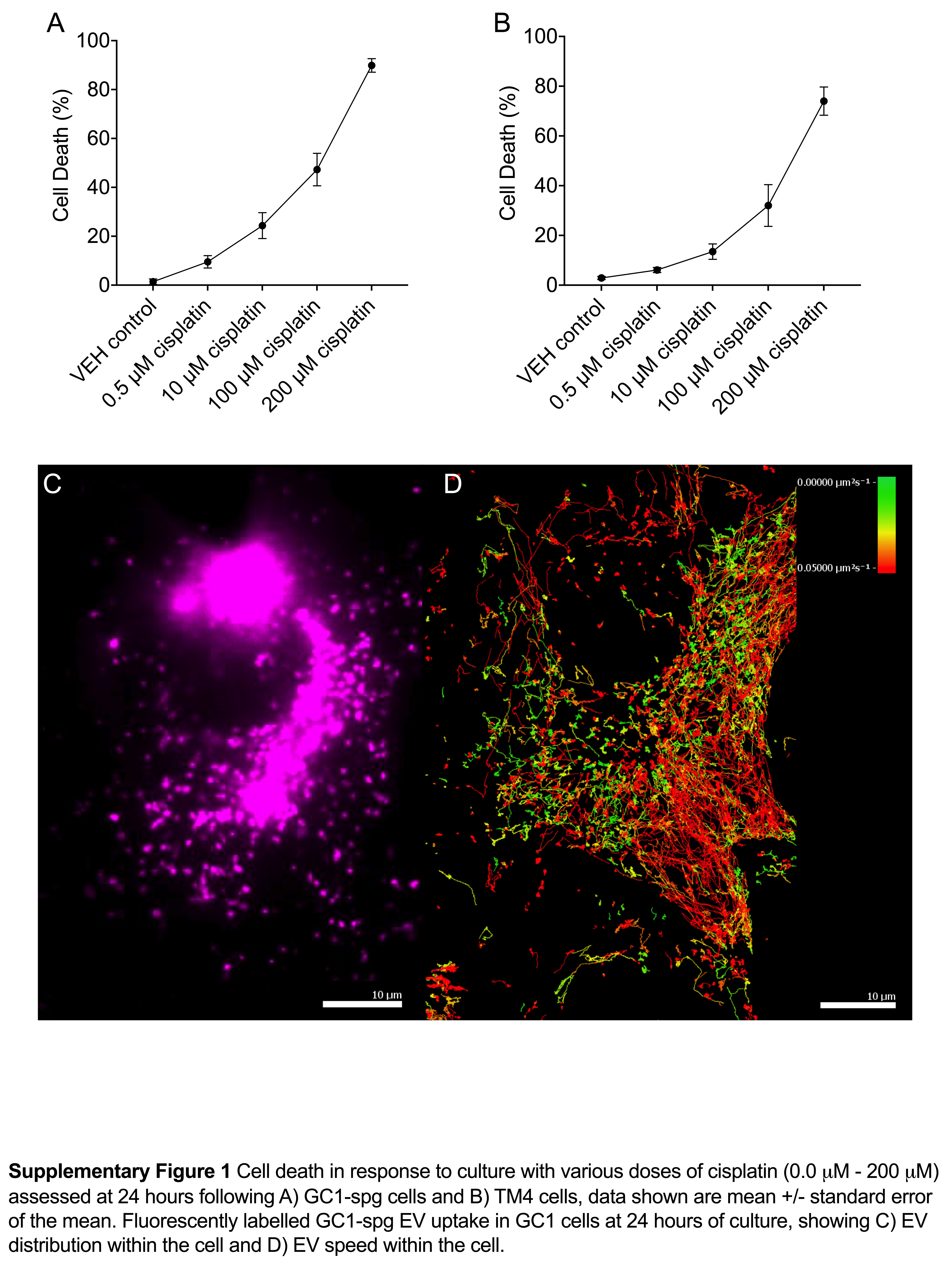

### Supplementary Figure 2

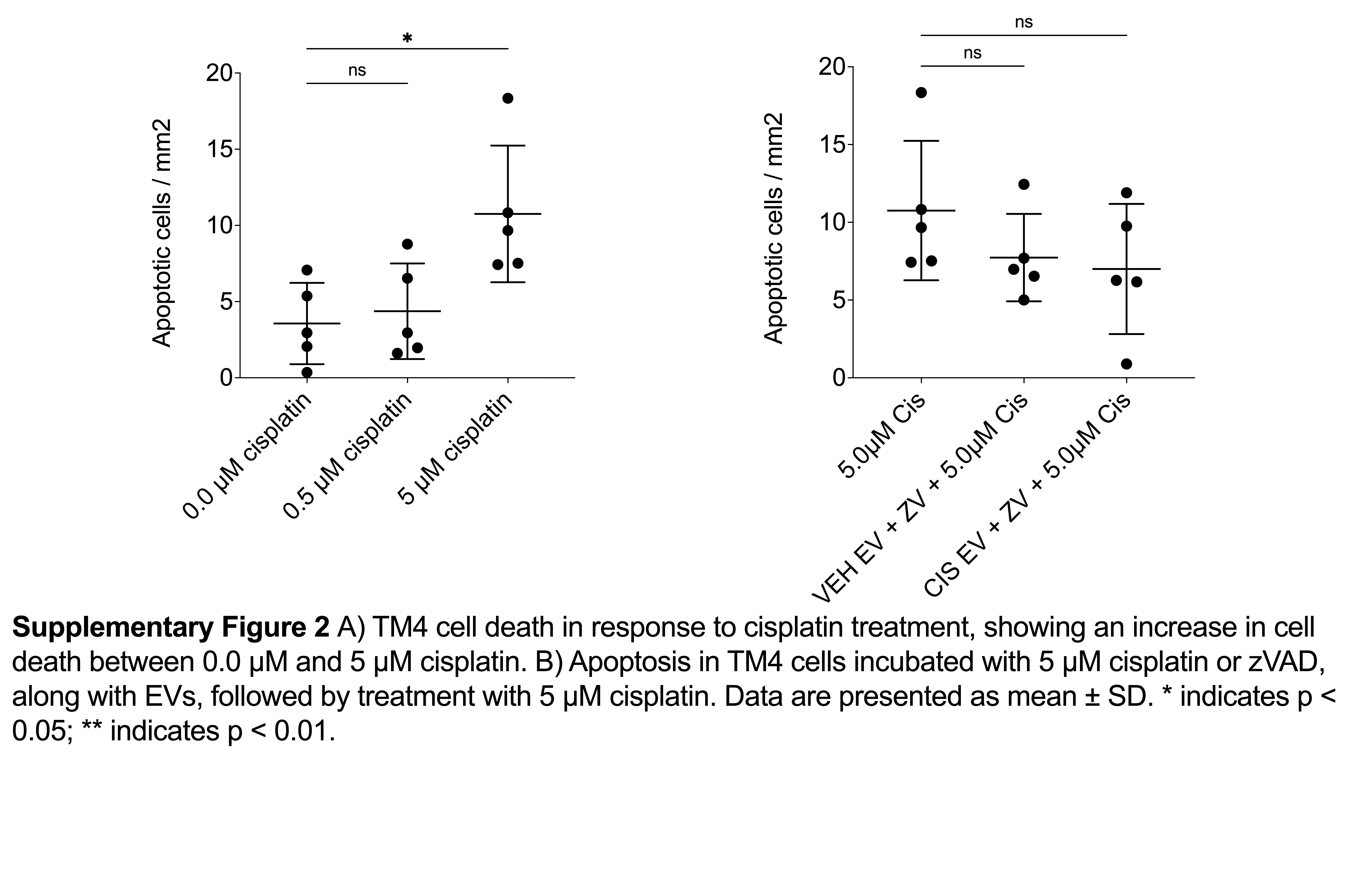
